# *Escherichia coli* has a Unique Transcriptional Program in Long-Term Stationary Phase

**DOI:** 10.1101/2020.04.28.067348

**Authors:** Karin E. Kram, Autumn Henderson, Steven E. Finkel

## Abstract

Microbes live in complex and consistently changing environments, but it is difficult to replicate this in the laboratory. *Escherichia coli* has been used as a model organism in experimental evolution studies for years; specifically, we and others have used it to study evolution in complex environments by incubating the cells into long-term stationary phase (LTSP) in rich media. In LTSP, cells experience a variety of stresses and changing conditions. While we have hypothesized that this experimental system is more similar to natural environments than some other lab conditions, we do not yet know how cells respond to this environment biochemically or physiologically. In this study, we begin to unravel the cells’ responses to this environment by characterizing the transcriptome of cells during LTSP. We found that cells in LTSP have a unique transcriptional program, and that several genes are uniquely upregulated or downregulated in this phase. Further, we identified two genes, *cspB* and *cspI*, which are most highly expressed in LTSP, even though these genes are primarily known to respond to cold-shock. When competed with wild-type cells, these genes are also important for survival during LTSP. These data allow us to compare biochemical responses to multiple environments and identify useful model systems, identify gene products that may play a role in survival in this complex environment, and identify novel functions of proteins.

**Importance:** Experimental evolution studies have elucidated evolutionary processes, but usually in chemically well-defined and/or constant environments. Using complex environments is important to begin to understand how evolution may occur in natural environments, such as soils or within a host. However, characterizing the stresses cells experience in these complex environments can be challenging. One way to approach this is by determining how cells biochemically acclimate to heterogenous environments. In this study we begin to characterize physiological changes by analyzing the transcriptome of cells in a dynamic complex environment. By characterizing the transcriptional profile of cells in long-term stationary phase, a heterogenous and stressful environment, we can begin to understand how cells physiologically and biochemically react to the laboratory environment, and how this compares to more natural conditions.

## Introduction

Experimental evolution studies of bacteria have revealed many insights into evolutionary processes (1–3). Many of these experiments have been performed in environments where only one factor is being experimentally manipulated, for instance starvation for one nutrient (4), heat shock (5), antibiotic stress (6), etc. However, in natural environments, cells are likely experiencing multiple, as well as varying, stresses. We and others have previously used long-term batch culture experimental evolution to explore how evolutionary processes work in a heterogenous environment (7–9). In these experiments, we allow cells to move through the entire life-cycle of *E. coli* in complex media, in order for cells to experience multiple types of stresses, which change throughout the incubation period.

During the *E. coli* life cycle in the laboratory, cells transition through the lag, log, and stationary phases then into death phase, where ∼99% of cells die, lyse, and release their cellular contents into the medium (10). This allows the remaining ∼1% of cells to enter long-term stationary phase (LTSP), where they can use the detritus of lysed cells as carbon and energy sources (10–12). The stresses of long-term stationary phase likely include high pH, high oxidative stress but low oxygen, and a lack of readily metabolized nutrients, however none of these are characterized completely (10, 13, 14). We have previously shown that cells with mutations that may help cope with these stresses are selected for during this phase, and that as populations continue further into LTSP, there is turnover of different mutant genotypes depending on the medium conditions at any given time (7, 9).

While we have hypothesized that this dynamism of genotypes indicates that different subpopulations of cells are growing and dying within LTSP cultures, we do not know if this phase consists of a collection of cells reflecting the four previous phases (lag, log, stationary, and/or death), or if LTSP consists of cells acting uniquely. We know that cells in lag, log, and stationary phases have unique transcriptional programs (15), but while expression changes due to genetic effects have been analyzed (16) to our knowledge, the transcriptional profile of LTSP in *E. coli* has not been studied. Elucidating the transcriptional profile of LTSP could help determine whether it is a phase in and of itself, or whether it is more an amalgam of cells experiencing the other phases.

In this study, we have analyzed the transcriptome of *E. coli* throughout its laboratory life cycle, including death phase and LTSP. We have determined that the expression of a small set of genes is uniquely regulated during LTSP, indicating that this phase does have its own transcriptional program. Further, we hypothesize that the genes which are only upregulated in LTSP compared to other phases may be important for survival during this phase. We identify three genes in the cold-shock protein family, *cspB, cspF*, and *cspI*, which are upregulated in LTSP compared to the other four phases. Mutants of two of these genes (*cspB* and *cspI*) affect survival in LTSP when competing with wild-type cells. These results further indicate that 1) we may be able to use these types of data to characterize the biochemical and physiological responses to long-term stationary phase stresses, and 2) that we may be able to identify novel functions for proteins which have only been characterized in earlier phases of the life cycle. Understanding how cells respond biochemically to LTSP may allow us to use standard laboratory conditions as models to better understand natural environments.

## Results

### The transcriptome of cells in LTSP can be distinguished from cells in other phases

In order to determine if there is a unique transcriptional program in LTSP, we incubated cells in triplicate into long-term stationary phase, extracted whole cell RNA at six time points throughout the *E. coli* life cycle, and performed RNA-sequencing (Fig. 1). Time points included log phase (4 hours), late-log phase (8 hours), stationary phase (24 hours), death phase (72 hours), and two time points in LTSP (144 and 192 hours). We sequenced rRNA-depleted RNA from each cell population, and analyzed at least 6 million reads per sample using HTSeq and DESeq2 (see methods section for more detailed explanation) (17, 18).

**Figure 1.**
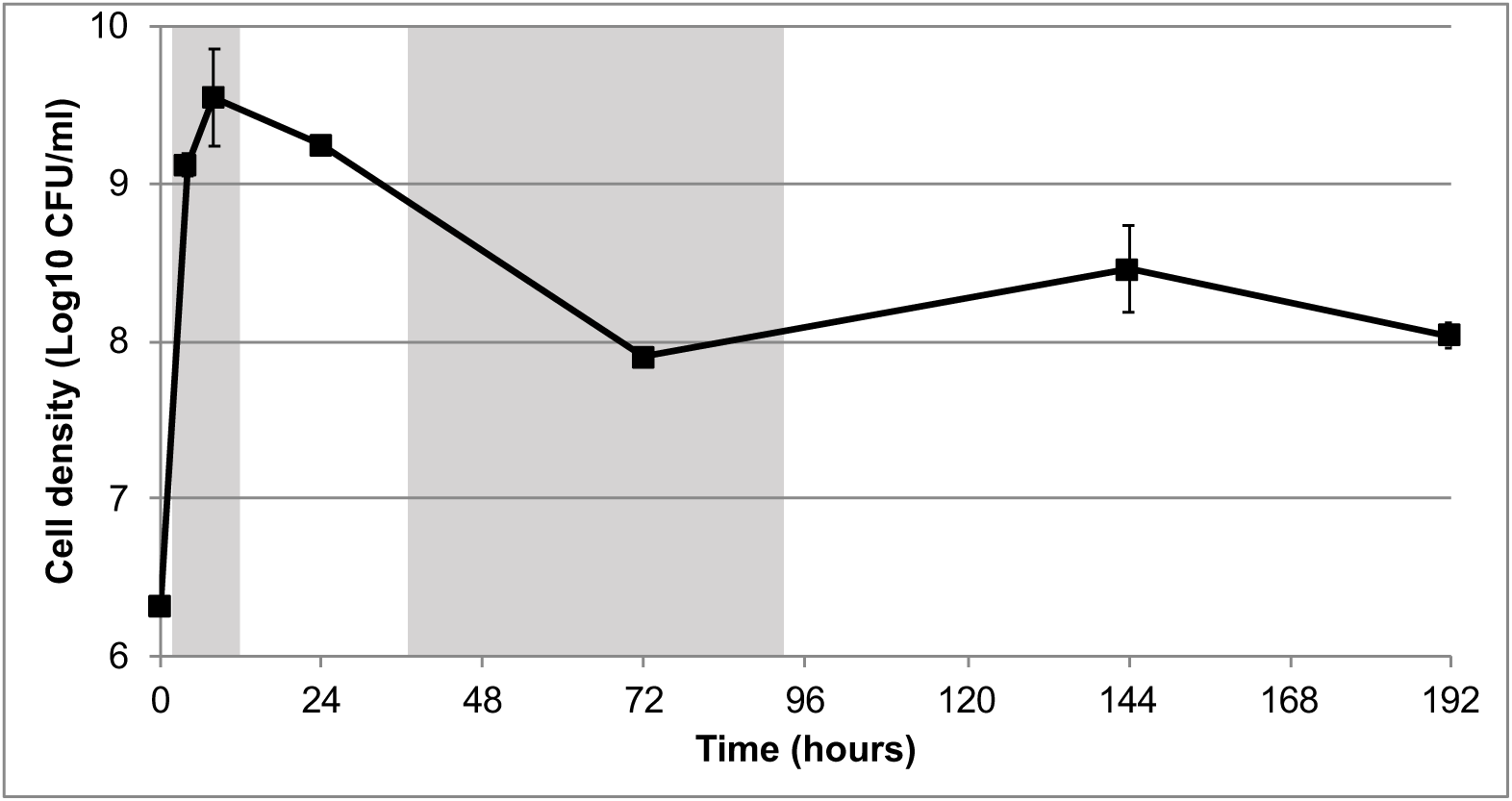
Average cell density during RNA-extraction time points. Three cultures were used for RNA extraction and for counting viable cells at 4, 8, 24, 72, 144, and 192 hours post inoculation. Points are an average of the three replicate cultures. Error bars represent standard deviation. In some cases error bars are so small that the point covers them. Shading represents a new phase in the *E. coli* life cycle (lag, log, stationary, death, long-term stationary).

Using the gene expression data, we performed a principal component analysis (PCA) to determine if there were features that distinguished the LTSP transcriptome from the other phases (Fig. 2). PCA ordinates the gene expression data across all 4437 genes included in the analysis to identify principal component axes (PCs) that capture the majority of the variance across all genes. We can plot each sample across these axes to observe how similar each sample is to the other sample with regard to all the expression data. We show comparisons of the samples among the first 3 PCs (Fig. 2), which together account for 73.4% of the variation in samples (PC1=31.9%, PC2=22.8%, and PC3=18.7%). Along PC1, we observed two clusters – one comprised of cells in log and late-log phase, and one with cells in stationary, death, and long-term stationary phase. Along PC2, we observed one cluster that includes cells in log, late-log, death phase, and LTSP, whereas cells in stationary phase are clustered on their own. And along PC3, we observed three clusters, one consisting of log and death phase cells, one with stationary phase and LTSP cells, and one that has late-log cells only. This observation shows that populations in LTSP have gene expression in common with the four other phases, and therefore supports the hypothesis that there would be sub-populations of cells in each of the first four phases during LTSP.

**Figure 2.**
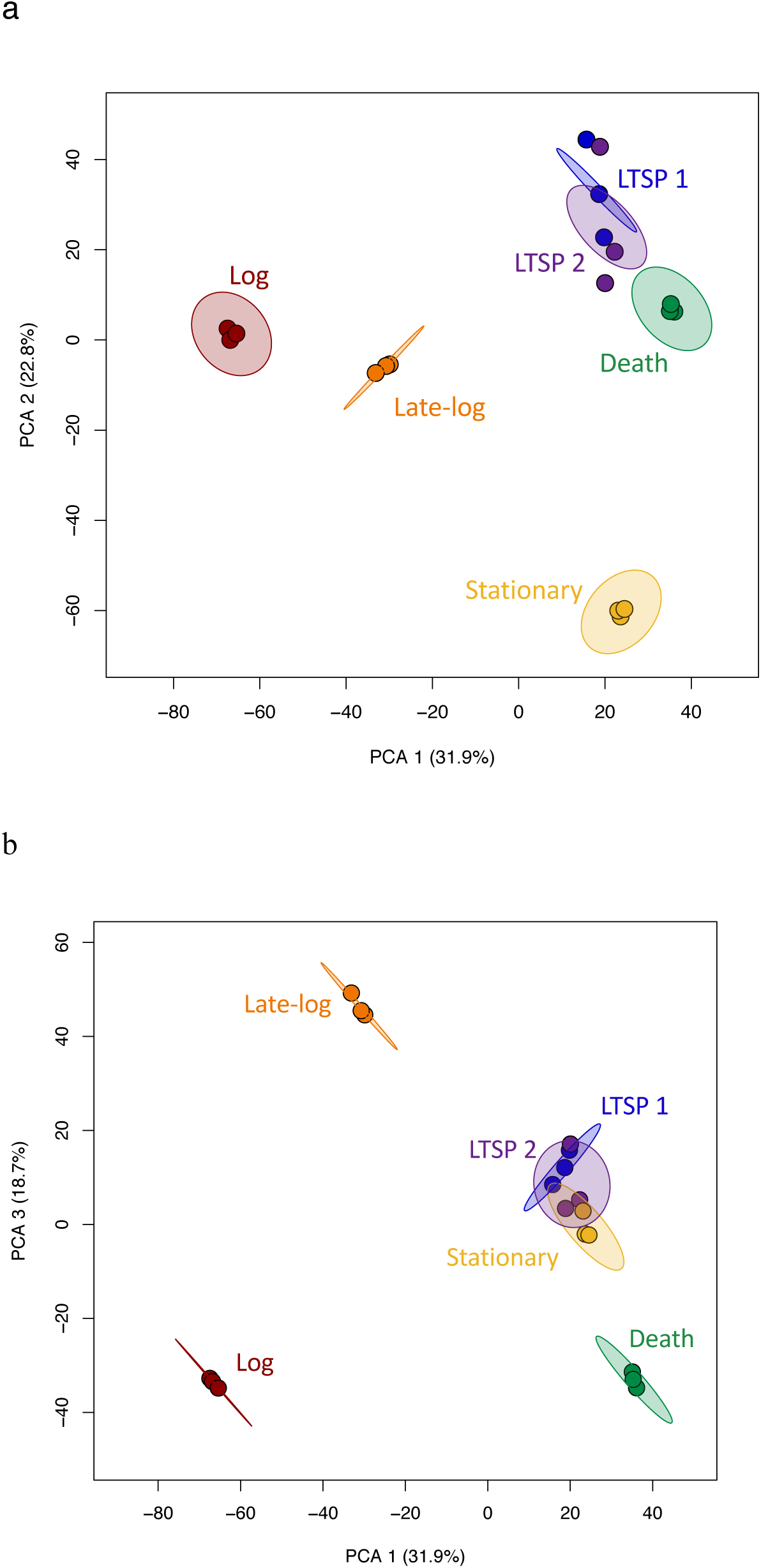
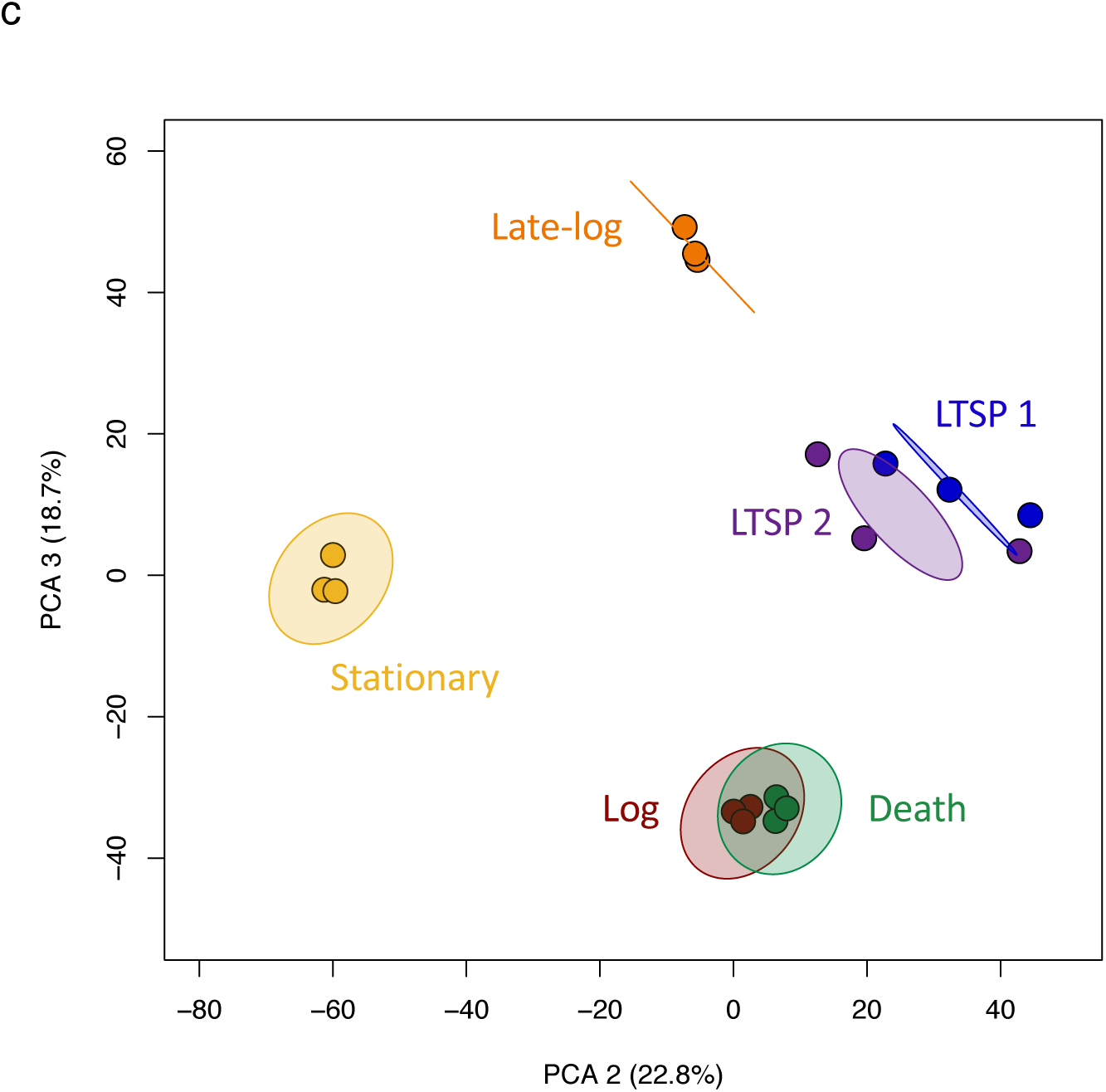
PCA of transcriptomes. Shaded circles represent 95% confidence intervals based on correlation matrices of the three replicates for each time point: 4h (red), 8h (orange), 24h (yellow), 72h (green), 144h (blue), 192h (purple). (a) PC1 vs. PC2 (b) PC1 vs. PC3 (c) PC2 vs. PC3. The PCA results suggest that cells in each phase have distinct gene expression patterns.

We also plotted a 95% confidence interval (CI) ellipse for each timepoint using a correlation matrix of our three replicates. By comparing overlap among ellipses for different timepoints, we can then determine how similar transcriptomes are across cells at different timepoints. These data indicate that cells mostly have unique transcriptional profiles in each stage of their life cycle (they stay within their own CI which does not overlap with another population’s CI), but there may be points throughout the life cycle when cells in one phase may also have subsets of gene expression in common with cells in other phases.

We noted that the 3 replicates for each time point before the cells reach LTSP cluster within their 95% CI, indicating that biological replicates respond very similarly to their environment even as it changes. Further we observed that populations in LTSP are not as similar to each other as those in other phases, as these replicates do not all appear within the 95% confidence interval. We also observed that the 95% CIs for each time point only overlap in the following cases: each LTSP time point overlaps with the other LTSP time point’s CI in each PC comparison (Fig. 2), the LTSP CIs overlap with stationary phase time points when comparing PC1 to PC3 (Fig. 2b), and log and death phase CIs overlap when comparing PC2 to PC3 (Fig. 2c).

The PC analysis indicates that cells harvested at the two LTSP time points are very similar to each other. In order to determine how similar the expression profiles were at these time points, we identified the differentially expressed (DE) genes between cultures incubated for 144 and 192 hours. Only 13 genes in 4 operons were determined to be significantly different (q value < 0.05, fold change > 2) between cells at these two time points (Table S1). Since this is a relatively small number compared to differences during other phases (for instance, 567 genes are significantly DE between log phase and 144h cells), we treated cells from the two LTSP time points as one group (LTSP) for the remainder of the analyses.

### Several genes are uniquely regulated in cells experiencing LTSP

While the PC analysis indicates that the cell population in LTSP is transcriptionally distinct from the other phases, it was not clear whether there were genes uniquely expressed in this phase, or whether this distinct profile was due to a mixture of cells in one of the other four stages.

In order to address this question, we compared the expression levels for each gene between LTSP and each of the four other phases. All statistically significant DE genes can be found in Table S2. Figure 3 shows a heatmap of each comparison containing all genes in the *E. coli* genome (Fig. 3). Of genes DE in LTSP from log phase, 755 are upregulated and 823 are downregulated during LTSP; of genes DE in LTSP from late-log phase 669 are upregulated and 463 are downregulated in LTSP; of genes DE in LTSP from stationary phase 632 are upregulated and 718 are downregulated in LTSP; and of genes DE in LTSP from death phase 311 are upregulated and 499 are downregulated in LTSP. In each comparison, with the exception of late-log phase, more genes are downregulated in LTSP than upregulated. Further, the number of upregulated genes decreases as populations approach LTSP.

**Figure 3.**
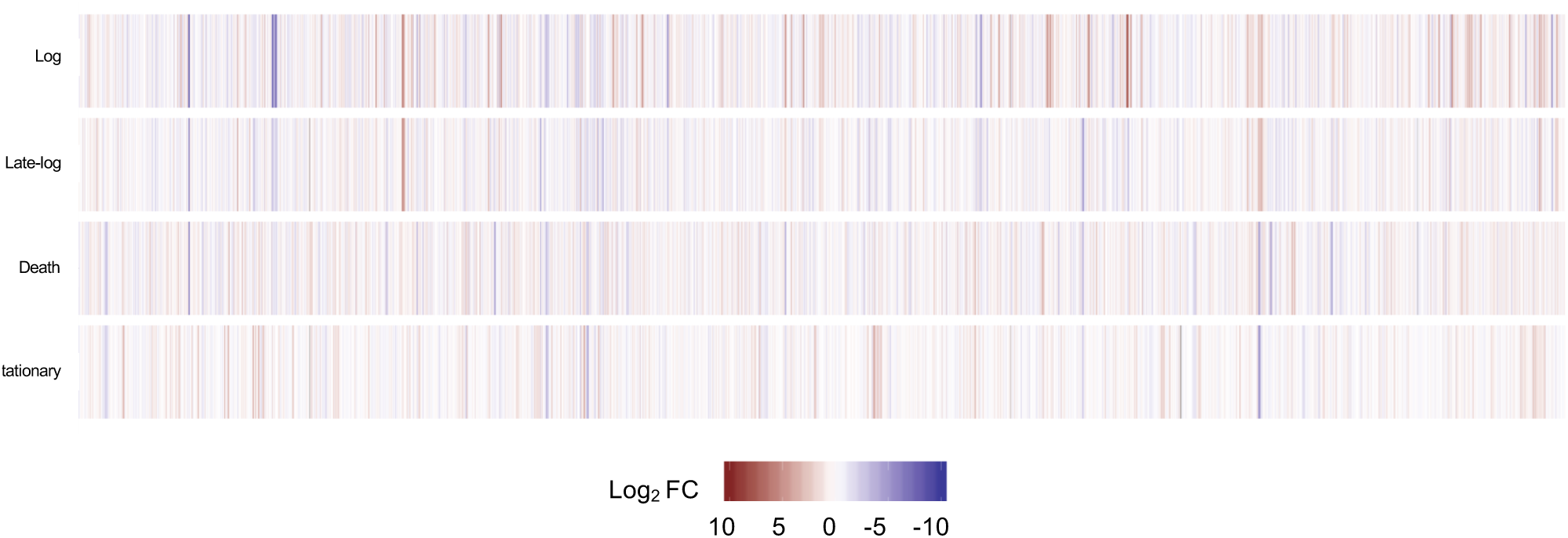
Heatmap of the expression difference between LTSP and other time points. Genes are along the x-axis in genome order.

While some genes are consistently DE from LTSP, there are many sets of genes that are DE only in one or two other phases. These data support the hypothesis that the population of cells in LTSP likely consists of cells experiencing one of the other phases, but that LTSP may also have a unique transcriptional program that some subset of cells are expressing.

In order to identify gene expression patterns unique to LTSP, we identified 38 genes that are significantly DE in LTSP compared to all other phases (Table 1); 32 of these genes are upregulated, and 6 of these genes are downregulated.

**Table 1.**
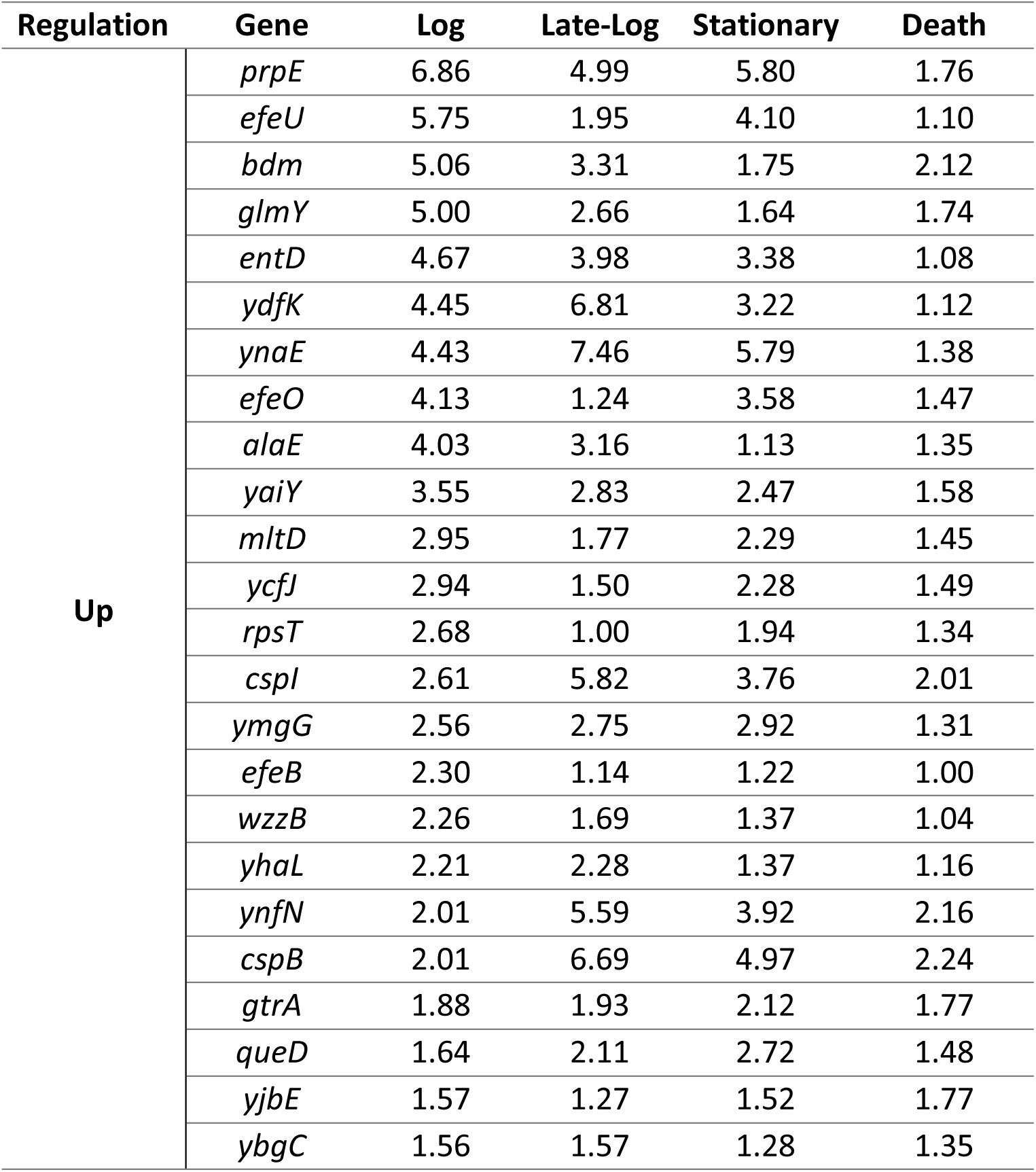
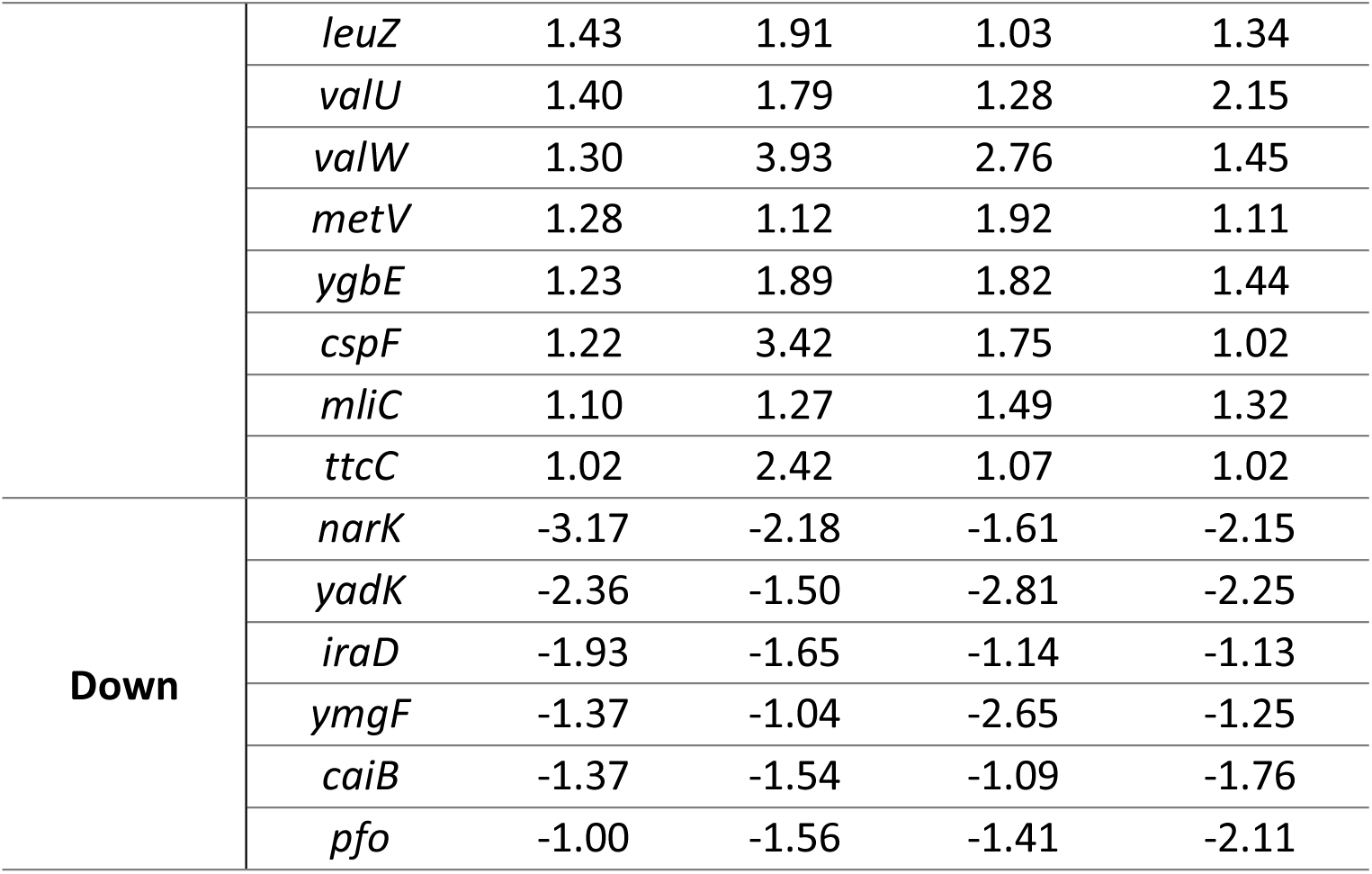
Log_2_ fold change of genes differentially expressed in LTSP compared to all other phases.

### Some genes uniquely expressed during LTSP affect survival during this phase

We hypothesize that genes which are expressed most highly in LTSP compared to other time points may play a role in cell survival. Interestingly, 3 genes identified as upregulated uniquely in LTSP were *csp* genes: *cspB, cspF*, and *cspI* (Table 1). *csp* genes were initially characterized as important for cells to physiologically adapt to cold temperatures, which cells in our cultures do not experience. Therefore, we were especially interested in this set of genes as they may be playing a role other than in response to cold shock in long-term batch cultures. The expression patterns of all 9 known *csp* genes during the incubation period is shown in Figure 4. We grouped these genes into 4 categories: (I) the genes mentioned above which are upregulated specifically during LTSP compared to all other phases (*cspB, cspF, cspI*), (II) genes which are highly expressed (normalized count > 1000) at all time points (*cspC, cspD, cspE*), (III) genes whose expression changes throughout the time course, but whose expression in LTSP is not different from all other phases (*cspA, cspG*), and (IV) one gene which shows low expression (normalized count < 15) at all time points (*cspH*). While *cspF* and *cspG* expression is at a similar level in LTSP, *cspF* expression is lower earlier in the life cycle, which is why statistically it should be placed in category I, whereas *cspG* expression starts higher earlier in the life cycle, which is why statistically it is grouped in category III (Fig. 4). Each *csp* gene is expressed from its own operon – none share promoters, although some may share regulatory regions – and are therefore likely regulated independently of one another.

**Figure 4.**
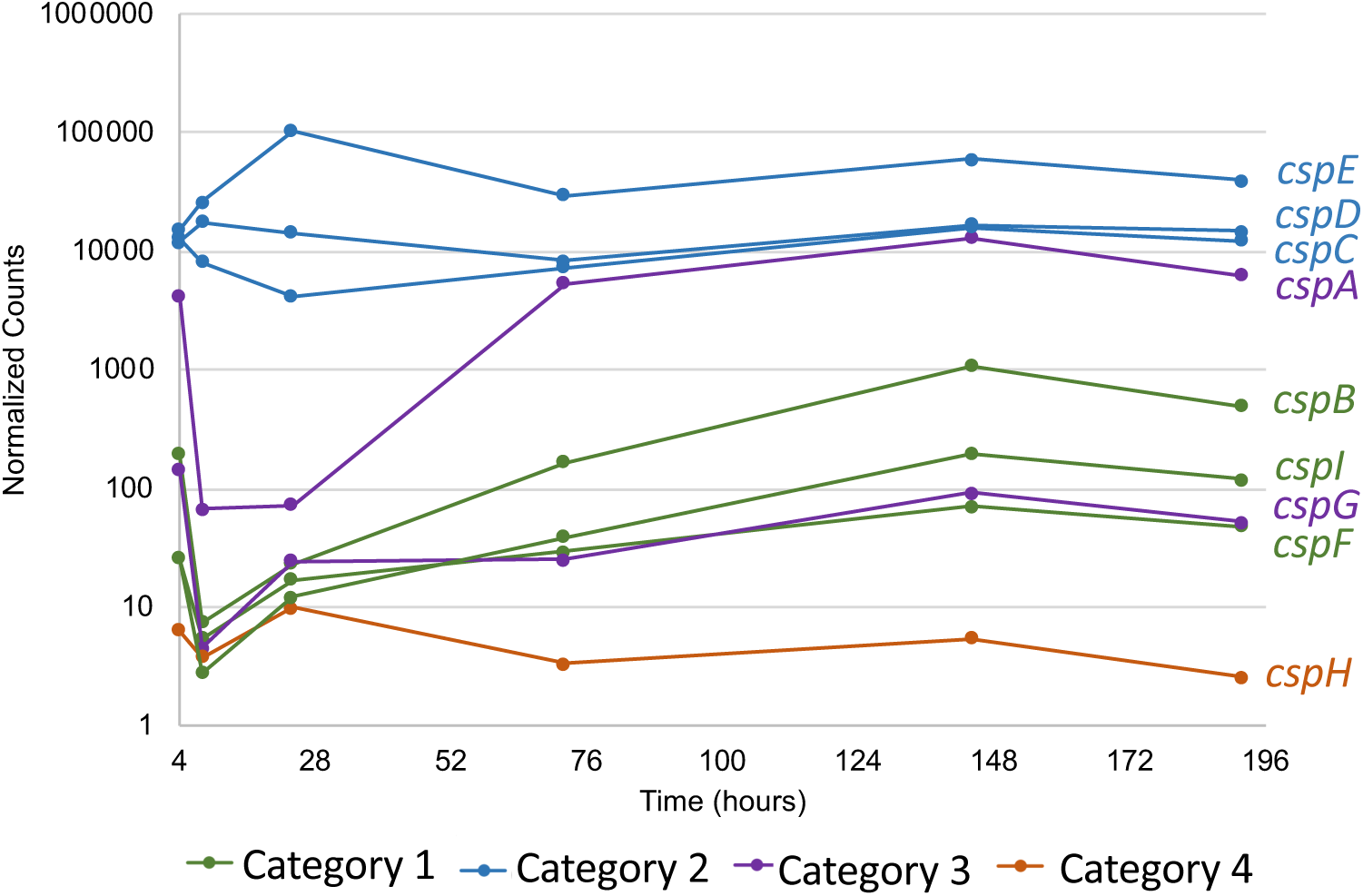
Expression pattern of *csp* genes. Normalized counts were output from DESeq2 analysis. Each point represents an average of the three cultures. We hypothesized that genes in category 1 would be important for survival in LTSP, genes in categories 2 and 3 may play a role in other phases of the life cycle, and the gene in category 4 would have no effect on cell growth or survival.

We predicted that all genes in category I would be important for survival in LTSP specifically, that genes in categories II and III may be important for growth or survival during other phases, and that genes in category IV would not affect cells under these culture conditions. To test these hypotheses, we deleted each gene from the PFM2 parental strain using P1 transduction from the Keio collection of gene knockouts (19), and incubated these cells into LTSP both alone in monoculture and in competition with wild-type cells. The data show that loss of genes in categories II, III, and IV have negligible, if any, effect on growth of *E. coli* when competed with wild-type cells in long-term cultures (Fig. 5). Loss of *cspC* or *cspE* (both category II) may give cells a slight (∼5-10 fold) advantage in early-LTSP, but the advantage is gone 48 hours later. Loss of the category 1 gene, *cspF* also has no effect (Fig. 5f).

**Figure 5.**
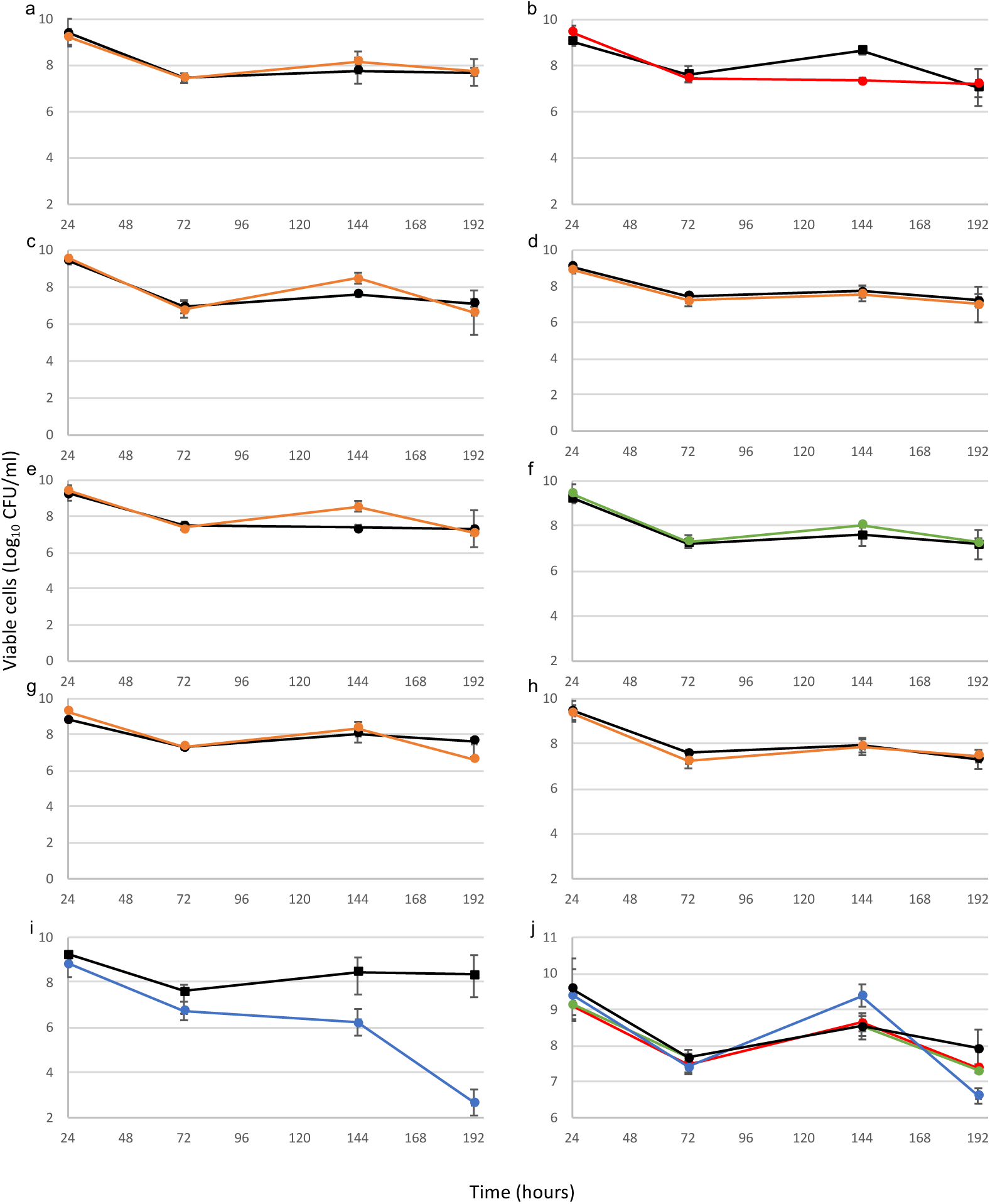
Competitions (a-i) and monocultures (j) of each *csp* mutant vs. wild-type cells. Wild-type cells are shown in black in each graph. Letters correspond with the csp gene that is missing in that mutant cell (i.e. (a) shows the competiton between the wild-type and *cspA*::kan^R^ strains), which the exception of (j), which shows monocultures of *cspB*::kan^R^ (red), *cspF*::kan^R^ (green), *cspI*::kan^R^ (blue) and WT (black). Note that (j) has a different y-axis scale. For (b), (f), (i), and (j) Points represent an average of three cultures. For (a), (c), (d), (e), (g), and (h), points represent an average of two cultures. Error bars represent standard deviation. In some cases error bars are so small that the point covers them.

However, loss of one of the other category 1 genes, *cspB* and *cspI*, does affect the cell’s ability to compete with the wild-type, to varying degrees (Figs. 5b and i). Cells mutant for *cspB* grow similarly to wild-type cells and experience a similar death phase, but then do not recover to the same level as wild-type cells at 144 hours (∼10-fold lower CFU/ml). In this case, wild-type cell density decreased over the next 48 hours, which brought them to the same level as the mutant cells (Fig. 5b). Cells lacking *cspI* again grow similarly to wild-type cells, but then have a slightly more severe death phase (∼10-fold lower CFU/ml at 72h), and never recover fully from death phase, leading to complete loss of viable mutant cells by 192h (Fig. 5i). Interestingly, when cells missing each of these genes are incubated in monoculture, without having to compete with wild-type cells, they do not show any defects in survival during death phase or LTSP (Fig. 5j).

## Discussion

We have determined that each phase in the *E. coli* life cycle has a unique transcriptional profile (Fig. 2). This has previously been observed for log phase and stationary phase, but to our knowledge this is the first indication that gene expression in death phase and LTSP is significantly different from other phases. Further, we have also shown that there are genes which are differentially expressed in LTSP as compared to all other phases, indicating that this phase has a unique transcriptional program, and is not only a combination of cells in other phases (although the populations are almost certainly also made up of cells in the other four phases, as well as those experiencing this unique transcriptional program).

At least two of the genes that are uniquely upregulated in LTSP, *cspB* and *cspI*, are important for survival in LTSP when competing against wild-type cells (Figs. 4 and 5). CspB and CspI are both cold-shock proteins, which were initially discovered as induced at low temperatures (20, 21). Generally, Csp proteins bind to RNA or ssDNA stabilizing it at low temperatures, which can also affect translational levels, although some Csp proteins, including CspB, may be chaperones for ssDNA as well (22, 23). Because cultures in these experiments are consistently incubated at 37**°**C, it is likely that these proteins are actually performing previously unidentified functions. Interestingly, both *cspB* and *cspI* are expressed in stationary phase even without cold-shock (20), further supporting the idea that these genes are likely being expressed in response to stresses other than cold-shock.

Other data indicate that these particular genes, as well as *csp* genes in general, may respond to other types of stresses. For instance, neither of these genes responded to cold stress in a pathogenic strain of *E. coli* (24). Further, Brandi et. al have shown that CspA, the originally identified cold-shock protein, can also be regulated by the global regulators Fis and H-NS, indicating that it likely responds to stresses other than cold shock (25). *cspC, cspD*, and *cspE* are regulated by growth arrest during diauxic shift, H_2_O_2_ stress, and the transition to stationary phase (26). CspC and CspE are important for virulence in Salmonella, again indicating that they likely respond to a signal other than cold-shock (27). Future studies will elucidate the roles of these and other genes during LTSP.

It is unsurprising that not all of the LTSP-specific genes identified play a specific role in LTSP – a similar finding was noted with transcriptome data in long-term cultures of *Rhodopseudomonas palustris* (28). However, data presented here indicate that, in fact, we can use expression patterns in LTSP to identify at least some genes that are important for survival in this environment and, further, possibly determine the growth state of cells within particular environments.

Determining which genes are expressed or repressed in lower nutrient environments, as well as which proteins are essential for survival, will allow us to characterize the biochemical and physiological responses to stress in LTSP. This will enable us to compare those biochemical responses to those in cells in natural environments, such as in a host or soil, to determine whether LTSP is a good model for these natural environments.

## Materials and Methods

### Bacterial strains, growth conditions, and viable cell counts

All strains in this study originate from strain PFM2, derived from the *E. coli* K-12 lineage strain, MG1655 (29) kanamycin resistant (kan^R^) and chloramphenicol resistant (cam^R^) derivatives or either mutant or “wild-type” cells, as appropriate. Before initiating any experiment, overnight cultures were inoculated from frozen stocks into 5 ml of Luria-Bertani (Lennox) medium (LB) (Difco) in 18-by 150-mm borosilicate test tubes, which were incubated with aeration in a TC-7 rolling drum (New Brunswick Scientific, Edison, NJ) at 37°C. We monitored cell growth and survival as described previously (13). Briefly, after inoculation at 1:1000 (vol:vol) dilution of cells from overnight growth frozen stocks into 5 ml of LB, we determined viable cell counts by plating serially dilutions of cultures on LB agar plates supplemented with kanamycin (50 μg/ml) or chloramphenicol (50 μg/ml), as appropriate.

### RNA preparation and sequencing

We inoculated all cultures for transcriptome analysis from a single PFM2 overnight culture. At each time point, we sacrificed 3 cultures to remove 1ml (4 hours), 0.5ml (8 hours), or 2ml (24, 72, 144, and 196 hours) of cells, and viable cell counts were determined by serial dilution as described above. RNAprotect reagent (Qiagen) was added per manufacturer’s instructions. Cells were pelleted and frozen at -80°C until total cellular RNA extraction was performed using the RNeasy Protect Bacteria Mini Kit (Qiagen). Total RNA was processed, assessed for quality, and sequenced by the USC Genomics Core, where ribosomal RNA was removed using the Ribo-zero Kit (Illumina, Inc.). 75-bp single end reads were generated from each sample on the Illumina HiSeq 2500 platform. We received an average of 7,882,901 reads per sample, with a range of ∼6.5 – 9.5 million reads per sample.

### Transcriptome analysis

Using a custom Python pipeline, we used TopHat2 to align the raw reads to the *E. coli* MG1655 genome (30), SAMtools to generate binary alignment files (BAM) (31), and HTSeq to calculate read counts per gene and per sample (18). An average of 89.4% of reads across all samples mapped to our reference genome (*Escherichia coli* K-12 MG1655), with a range of ∼66-96%. All ribosomal RNA gene reads, as well as any genes that had no expression, were removed from the dataset. We used the HTSeq output files as input to analyze gene expression levels using DESeq2 (17), with comparisons between samples made based on the time of incubation. Once we determined that expression differences between the 144h and 192h were minimal, we marked these data points as the same time (“LTSP”) in our dataframe. We identified differentially expressed genes using pairwise comparisons between time points. We considered genes to be differentially expressed between cells in LTSP and cells at other time-points if the log_2_ fold change was greater than or equal to 1, and if the q value was less than or equal to 0.05. We summarized the data using principal component analysis (PCA) using the log scale values of the normalized counts of each gene using the prcomp function using centering and scaling in base R. We added 95% confidence intervals by calculating correlation matrices for each the samples in each timepoint, and then adding these intervals to our plot using the polygon function in the ellipse package in R (32). We also created a heatmap in ggplot2 (33) comparing the log_2_ fold change between the average normalized counts for each gene in LTSP versus each other time point using ggplot2 (33).

### Mutagenesis of *csp* genes

We constructed in-frame knock-out mutations of each *csp* gene individually (*cspA* to *cspI*) to create nine mutant strains using P1 transduction from strains in the Keio collection (19). Briefly, we inoculated overnight cultures from a frozen stock in the Keio collection. The following day, we inoculated cultures at a 1:100 dilution into fresh medium and allowed them to grow to mid-log phase. Lysates were made from the donor cells using P1 stock. We then used the lysates to transduce wild-type PFM2, and selected for mutants on LB agar plates supplemented with kanamycin. We confirmed gene replacement using PCR.

### Competitions and monocultures with wild-type and mutant strains

For monocultures, we inoculated cultures as described above. We also performed competitions between mutant strains and parental strains. Parental strains were marked with a chloramphenicol cassette replacing *lacZ* (*lacZ*::cam^R^) (7) and mutant strains were marked with a kanamycin cassette replacing the gene of interest (*cspA*::kan^R^ to *cspI*::kan^R^). We inoculated wild-type (*lacZ*::cam^R^) and each of the mutant cells (*csp*\*::kan^R^) from frozen stocks as described above, and overnight cultures were each co-inoculated 1:1000 (vol:vol) into LB broth in test tubes. We assayed both monoculture and competition populations for viable counts throughout the experiment (0, 4, 8, 24, 72, 144, and 192 hours after inoculation) as described above.

### Data availability

Both count files derived from HTseq and raw sequence reads have been deposited in the GEO database and are available.

## Acknowledgements

We would like to thank Dr. Lacey L. Westphal for help with data analysis and Professor Daniel Stoebel, Professor Sonal Singhal and members of the Kram lab for helpful discussion of the project and manuscript.

This work was supported in part by U.S. Army Research Office grants W911NF1010444 and W911NF1210321 (to S.E.F.) and by National Science Foundation RUI #1715006 (to K.E.K). K.E.K. was also partially supported by the CSUDH Research, Scholarship, and Creative Activity award, the CSUDH Norris Foundation Award, and the CSUDH Emeritus Faculty Fund Award. The funders had no role in study design, data collection and interpretation, or the decision to submit the work for publication.

---

**Supplemental Table 1**. Differentially expressed genes between LTSP time points.

**Supplemental Table 2**. Differentially expressed genes between LTSP and at least one other phase.

## References

1. Lenski RE, Wiser MJ, Ribeck N, Blount ZD, Maddamsetti R, Burmeister AR, Baird EJ, Bundy J. 2015. Sustained fitness gains and variability in fitness trajectories in the long-term evolution experiment with *Escherichia coli*. Proceedings of the Royal Society B 282:201522292.

2. Maharjan RP, Liu B, Feng L, Ferenci T, Wang L. 2015. Simple phenotypic sweeps hide complex genetic changes in populations. Genome Biology and Evolution 7:531–544.

3. Hindré T, Knibbe C, Beslon G, Schneider D. 2012. New insights into bacterial adaptation through *in vivo* and *in silico* experimental evolution. Nature Reviews Microbiology 10:352–365.

4. Blount ZD, Borland CZ, Lenski RE. 2008. Historical contingency and the evolution of a key innovation in an experimental population of *Escherichia coli*. Proceedings of the National Academy of Sciences 105:7899–906.

5. Rodŕiguez-Verdugo A, Tenaillon O, Gaut BS. 2016. First-step mutations during adaptation restore the expression of hundreds of genes. Molecular Biology and Evolution 33:25–39.

6. Toprak E, Veres A, Michel J-B, Chait R, Hartl DL, Kishony R. 2011. Evolutionary paths to antibiotic resistance under dynamically sustained drug selection. Nature Genetics 44:101–105.

7. Westphal LL, Lau J, Negro Z, Moreno IJ, Ismail Mohammed W, Lee H, Tang H, Finkel SE, Kram KE. 2018. Adaptation of *Escherichia coli* to long-term batch culture in various rich media. Research in Microbiology. 169:145–156.

8. Chib S, Ali F, Seshasayee ASN. 2017. Genomewide mutational diversity in *Escherichia coli* population evolving in prolonged stationary phase. mSphere 2:e00059–17.

9. Kram KE, Geiger C, Ismail WM, Lee H, Tang H, Foster PL, Finkel SE. 2017. Adaptation of *Escherichia coli* to long-term serial passage in complex medium: Evidence of parallel evolution. mSystems 2:e00192–16.

10. Finkel SE. 2006. Long-term survival during stationary phase: evolution and the GASP phenotype. Nature Rev Microbiol 4:113–120.

11. Takano S, Pawlowska BJ, Gudel I, Yomo T, Tsuru S. 2017. Density-dependent recycling promotes the long-term survival of bacterial populations during periods of starvation. mBio 8:e02336–16.

12. Schink SJ, Biselli E, Ammar C, Gerland U. 2019. Death rate of *E. coli* during starvation is set by maintenance cost and biomass recycling. Cell Systems 9:64–73.

13. Kram KE, Finkel SE. 2015. Rich medium composition affects *Escherichia coli* survival, glycation, and mutation frequency during long-term batch culture. Applied and Environmental Microbiology 81:4442–4450.

14. Kram KE, Finkel SE. 2014. Culture volume and vessel affect long-term survival, mutation frequency, and oxidative stress in *E. coli*. Applied Environmental Microbiology 80:1732–1738.

15. Chang DE, Smalley DJ, Conway T. 2002. Gene expression profiling of Escherichia coli growth transitions: An expanded stringent response model. Molecular Microbiology 45:289–306.

16. Arunasri K, Adil M, Khan PA, Shivaji S. 2014. Global gene expression analysis of long-term stationary phase effects in *E. coli* K12 MG1655. PLoS ONE 9:e96701.

17. Love MI, Huber W, Anders S. 2014. Moderated estimation of fold change and dispersion for RNA-seq data with DESeq2. Genome Biology 15:550.

18. Anders S, Pyl PT, Huber W. 2015. HTSeq--a Python framework to work with high-throughput sequencing data. Bioinformatics 31:166–169.

19. Baba T, Ara T, Hasegawa M, Takai Y, Okumura Y, Baba M, Datsenko KA, Tomita M, Wanner BL, Mori H. 2006. Construction of *Escherichia coli* K-12 inframe, single-gene knockout mutants: the Keio collection. Molecular Systems Biology 2:2006.0008-undefined.

20. Czapski TR, Trun N. 2014. Expression of *csp* genes in *E. coli* K-12 in defined rich and defined minimal media during normal growth, and after cold-shock. Gene 547:91–97.

21. Wang N, Yamanaka K, Inouye M. 1999. CspI, the ninth member of the CspA family of *Escherichia coli*, is induced upon cold shock. Journal of Bacteriology 181:1603–1609.

22. Yamanaka K, Fang L, Inouye M. 1998. The CspA family in *Escherichia coli*: multiple gene duplication for stress adaptation. Molecular Microbiology 27:247–255.

23. Yu T, Keto-Timonen R, Jiang X, Virtanen JP, Korkeala H. 2019. Insights into the phylogeny and evolution of cold shock proteins: From enteropathogenic *Yersinia* and *Escherichia coli* to eubacteria. International Journal of Molecular Sciences 20:4059.

24. Li Y, Zhou D, Hu S, Xiao X, Yu Y, Li X. 2018. Transcriptomic analysis by RNA-seq of *Escherichia coli* O157:H7 response to prolonged cold stress. LWT - Food Science and Technology 97:17–24.

25. Brandi A, Giangrossi M, Giuliodori AM, Falconi M. 2016. An interplay among FIS, H-NS, and guanosine tetraphosphate modulates transcription of the *Escherichia coli cspA* gene under physiological growth conditions. Frontiers in Molecular Biosciences 3:19.

26. Chang D-E, Smalley DJ, Conway T. 2002. Gene expression profiling of Escherichia coli growth transitions: an expanded stringent response model. Molecular Microbiology 45:289–306.

27. Michaux C, Holmqvist E, Vasicek E, Sharan M, Barquist L, Westermann AJ, Gunn JS, Vogel J. 2017. RNA target profiles direct the discovery of virulence functions for the cold-shock proteins CspC and CspE. Proceedings of the National Academy of Sciences 114:6824–6829.

28. Pechter KB, Yin L, Oda Y, Gallagher L, Yang J, Manoli C, Harwood CS. 2017. Molecular basis of bacterial longevity. mBio 8:e01726–17.

29. Foster PL, Lee H, Popodi E, Tang H. 2015. Determinants of spontaneous mutation in the bacterium *Escherichia coli* as revealed by whole-genome sequencing. Proceedings of the National Academy of Sciences 112:E5990–E5999.

30. Kim D, Pertea G, Trapnell C, Pimentel H, Kelley R, Salzberg SL. 2013. TopHat2: accurate alignment of transcriptomes in the presence of insertions, deletions and gene fusions. Genome Biology 14:R36.

31. Li H, Handsaker B, Wysoker A, Fennell T, Ruan J, Homer N, Marth G, Abecasis G, Durbin R. 2009. The Sequence Alignment/Map format and SAMtools. Bioinformatics 25:2078–2079.

32. Murdoch DJ, Chow ED. 1996. A graphical display of large correlation matrices. The American Statistician 50:178–180.

33. Wickham H. 2016. ggplot2: Elegant graphics for data analysis. Springer-Verlag New York.

